# Auditory Aversive Generalization Learning Prompts Threat-Specific Changes in Alpha-Band Activity

**DOI:** 10.1101/2023.12.04.569971

**Authors:** Andrew H. Farkas, Richard T. Ward, Faith E. Gilbert, Jourdan Pouliot, Payton Chiasson, Skylar McIlvanie, Caitlin Traiser, Kierstin Riels, Ryan Mears, Andreas Keil

## Abstract

Pairing a neutral stimulus with aversive outcomes prompts neurophysiological and autonomic changes in response to the conditioned stimulus (CS+), compared to cues that signal safety (CS-). One of these changes—selective amplitude reduction of parietal alpha-band oscillations— has been reliably linked to processing of visual CS+. It is however unclear to what extent auditory conditioned cues prompt similar changes, how these changes evolve as learning progresses, and how alpha reduction in the auditory domain generalizes to similar stimuli. To address these questions, fifty-five participants listened to three sine wave tones, with either the highest or lowest pitch (CS+) being associated with a noxious white noise burst. A threat specific (CS+) reduction in occipital-parietal alpha-band power was observed similar to changes expected for visual stimuli. No evidence for aversive generalization to the tone most similar to the CS+ was observed in terms of alpha-band power changes, aversiveness ratings, or pupil dilation. By-trial analyses found that selective alpha-band changes continued to increase as aversive conditioning continued, beyond when participants reported awareness of the contingencies. The results support a theoretical model in which selective alpha power represents a cross-modal index of continuous aversive learning, accompanied by sustained sensory discrimination of conditioned threat from safety cues.

## Introduction

It has been well established that cues predicting aversive outcomes capture and hold access to limited capacity systems in the human brain, including systems mediating sensation, perception, and attention (Li & Keil, 2023). The neurophysiological processes involved in acquiring this privileged access are less understood, including how cues in different sensory modalities acquire sensitivity to conditioned threat. A growing body of work has consistently shown that pairing a visual cue with an aversive outcome over time prompts a greater power reduction in posterior alpha-band (8 – 13 Hz) oscillations compared to unpaired cues (Friedl & Keil, 2020; Panitz et al., 2019). Additionally, unpaired visual cues that are perceptually similar to the conditioned stimulus also prompt alpha-band power reduction in the same parieto-occipital areas, but to a lesser degree, creating a Gaussian pattern of effects (Friedl & Keil, 2021; Yin et al., 2020). These changes have been taken as evidence for widespread cortical disinhibition, consistent with concepts such as arousal or attention (Bacigalupo & Luck, 2022; Li & Keil, 2023). It is however unclear how auditory aversive conditioning affects alpha-band power. Neutral tones that are paired with aversive white noise bursts modulate the late positive potential (LPP) component of the event-related potential (ERP), similar to visual cue paradigms (Pavlov & Kotchoubey, 2019). The LPP in turn tends to mirror alpha-power changes during emotional scene perception (De Cesarei & Codispoti, 2011; Ferrari et al., 2020). In line with the notion that alpha power reduction may be a cross-modal index of aversive learning, Hartmann et al., (2012) found lowered alpha power in auditory cortex when participants were instructed that a tone cue would be followed by a noxious noise. However, the extent and temporal dynamics of alpha-power changes during associative learning are still to be established. Specifically, it remains unclear if auditory aversive conditioning results in selective, stimulus-specific alpha-band power reduction, and if so, which cortical areas contribute to these effects. Furthermore, it is unclear if auditory conditioning prompts generalization across similar tone cues—a hallmark of visual associative learning.

To address these questions, alpha-band power in human dense-array EEG was examined in a differential auditory conditioning paradigm in which the highest or lowest pitch of three tones were paired with a noxious white noise burst (Figure 1). To establish the extent to which learning took place, self-reports of valence and arousal as well as pupil dilation were collected in addition to EEG alpha power. In differential Pavlovian conditioning, an initially neutral stimulus (the CS+) such as a soft tone or visual pattern is repeatedly paired with an unconditioned stimulus (US) such as an electric shock or noxious noise. Other stimuli (the CS-) are never paired with the US. When multiple CS- are used, it is possible to examine response tuning along a gradient that is defined by perceptual similarity with the CS+. In this type of experiment, the CS- are then referred to as generalization stimuli (GS). In the present study, three tones were initially presented to participants without any aversive US pairing (the Habituation phase). Then, either the lowest or highest pitch was paired with a noxious US white noise (the Acquisition phase). Self-reports for each cue were collected at early and late periods of both the Habituation and Acquisition phases, while pupil dilation was recorded simultaneously with the EEG.

**Figure 1.**
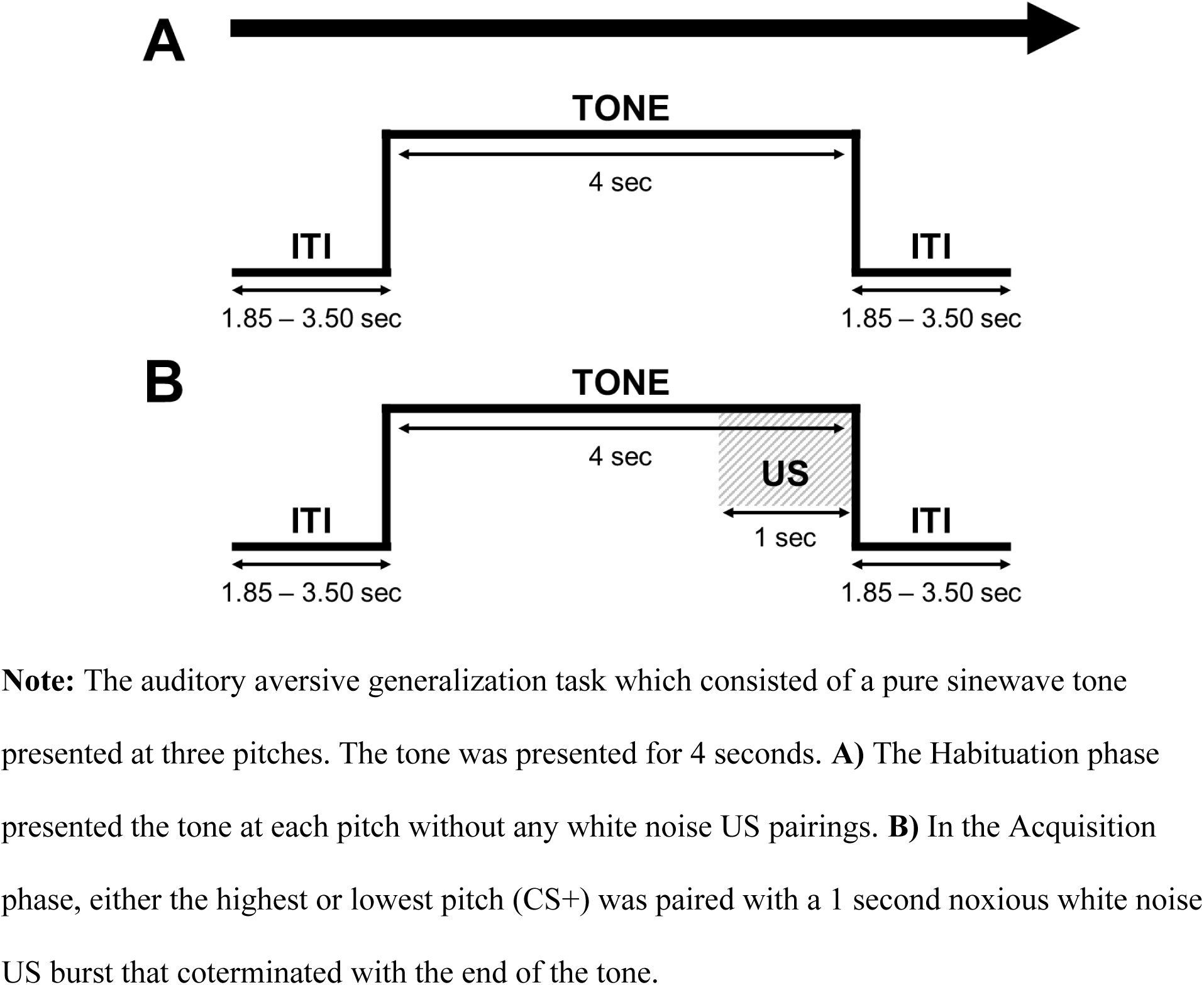
Auditory Aversive Generalization Task.

To focus and simplify the analyses, the results were interpreted through two a-priori models representing the most likely patterns of cue reactivity. As previously mentioned, occipital-parietal alpha-power changes are known to generalize to cues that are visually similar to CS+ (Friedl & Keil, 2021; Yin et al., 2020). Accordingly, it was hypothesized that threat-related changes in parietal alpha oscillations would generalize across a gradient of auditory stimuli varying in pitch. Another possibility is an all-or-nothing pattern in which there is only reactivity to the CS+. To represent these possible reactivity patterns, the models were specified by assigning a single weight-value for each cue representing the expected increase or decrease in reactivity. These weights were then used in planned comparison analyses, within the framework of the general linear model, also known as F-contrasts (Rosenthal & Rosnow, 1985). Both models predict an increase for the CS+ with a positive weight. The models differ in their expectations for the next closest tone pitch (GS1) and the least similar tone (GS2). The *All-or-Nothing* model predicts no change for GS1 and GS2 (weights: 2, −1, −1), whereas the *Generalization* model predicts an increase for GS1 and a reduction for GS2 (weights: 1, 0.75, and –1.75). The fits between the data and each of the two a-priori models can be thought of as competing alternative hypotheses of All-or-Nothing versus Generalization patterns of reactivity.

## Methods

The data presented in this report are part of a larger study design and set of hypotheses that were preregistered prior to data collection (https://osf.io/e26ad). The current study concerns findings regarding alpha-band power changes over an auditory stimulus generalization gradient across participants, focusing on within-subject effects.

### Participants

Sixty-four participants were recruited using flyers, existing databases, and online advertisements. The participants received class credit or were paid $20 USD per hour for completion of the study. All procedures were approved by the University of Florida institutional review board, and participants gave informed consent prior to participating in accordance with the Declaration of Helsinki. All participants were 18 years of age or older, had normal or corrected-to-normal vision, and reported no personal or family history of seizures. Nine participants were excluded from this analysis due to withdrawal (*n* = 5), or having over 50% of EEG trial data contaminated by artifacts (*n* = 4). This resulted in a total of 55 remaining participants (36 Female; *M_age_* = 20.34, *SE_age_* = 0.35) used for data analyses of ratings and alpha-band power. For the secondary analyses of pupil dilation, 20 participants from the EEG sample were removed due to unusable pupil recordings leaving 35 participants for the pupil dilation analyses (24 Female; *M_age_* = 20.21, *SE_age_* = 0.49). Pupil data were considered unusable when the majority of trials did not meet trial inclusion criteria described below.

Participants completed several other measures related to defensive engagement in aversive learning that are not reported here as the present focus is on overall reactivity not influenced by individual differences. These measures included questionnaires related to Misophonia, fear, anxiety, and depression symptoms. Individuals reporting elevated Misophonia symptoms (i.e., scores above 19 on the Misophonia Symptom Scale; (Wu et al., 2014)) were not recruited into this sample. Loudness discomfort levels were also acquired as an index of hyperacusis. No participants in the primary sample reported elevated loudness discomfort ratings (*M* = 23.90, *SE* = 2.22, on a scale from 5 to 50); the same was true for the sub-sample which was used for pupil analyses (*M* = 19.30, *SE* = 2.50).

### Materials & Procedures

#### Ratings of Affect Using the Self-Assessment Manikin

Ratings for arousal and valence were collected using the Self-Assessment Manikin (SAM; (Bradley & Lang, 1994)) during the 10^th^ (early) and 90^th^ (late) trials of the Habituation and Acquisition phases of the experiment. After each tone at this specific point of the experiment, participants self-reported their experienced arousal and valence. Each dimension was presented with a set of five manikins ranging from completely calm/pleasant (i.e., low arousal or low valence scores) to aroused/unpleasant (i.e., high arousal or high valence scores). While viewing the manikins, participants reported how the previous tone made them feel by moving a monitor cursor with a mouse and clicking to make their rating. Responses were recorded as the pixel location along the horizontal axis of the SAM which ranged from 1 to 1,920 pixels.

#### Auditory Stimuli

The auditory stimuli were three sinewave tones (sampled at 22,000 Hz) that were four seconds in duration. A cosine square window (20 points ramp-on and ramp-off) was applied at the beginning and end of the auditory presentation to minimize onset and offset sound spikes. Each sinewave tone had a specific frequency (i.e., 320, 541, or 914 Hz) based on an exponential function, creating three pitch conditions (i.e., CS+, GS1, and GS2). The number of tones and the pitch frequencies were chosen based on pilot studies, such that the majority of participants could discriminate the tones. Additionally, tones were multiplied by a 41.2 Hz cosine envelope to evoke auditory steady-state potentials, used in a separate set of hypotheses and analyses not reported here (see preregistration for additional information). Each tone was normalized based on their respective frequency’s amplitude. Finally, the loudness levels of each tone were reduced to 20% of their amplitude, resulting in loudness levels of 70 dBA for each tone stimulus, assessed with an audiometer.

During the acquisition phase, a 1-second 91 dBA white noise was played during CS+ trials, serving as the US, and accompanying the CS+ during the final second of the four second tone duration. The US duration and onset were selected based on previous research suggesting larger conditioning effects for durations over 500 ms (Sperl et al., 2016), and when a CS+ and US co-terminate following prolonged overlapping intervals (Kamin, 1956). Levels were controlled with an audiometer between sessions. The white noise also included a short cosine-square ramp (5 points) at the beginning and end of this auditory presentation. The white noise was paired with either the 320 Hz or 914 Hz pitch, counterbalanced across participants. Thus, the paired pitch served as CS+ (100% reinforcement rate), with the other frequencies considered as GS1 or GS2, depending on their distance in frequency from the CS+ pitch. All auditory stimuli were presented through two Behringer Studio 50 speakers arranged symmetrically behind each participant at ear level, approximately 30 cm in distance from the participant.

#### Auditory Aversive Generalization Task

The auditory aversive generalization task featured 240 total trials split into two 120 trial phases, Habituation (Figure 1A) and Acquisition (Figure 1B). In accordance with ethical standards for this paradigm, this was followed by an extinction phase in which trials were presented without a US pairing. The data for the extinction phase was excluded because the paradigm was designed for investigating continued associative learning during acquisition. Thus, the extinction phase did not include enough trials for interpretable results. For each trial, one of the three tones (i.e., CS+, GS1, GS2) was played for 4 seconds while participants viewed a white fixation dot on a monitor occupying 0.5° of visual angle. The white noise US was only presented during the Acquisition phase and was paired with the CS+ pitch (i.e., 320 or 914 Hz) at 100% reinforcement rate. The SAM ratings of arousal and valence for each pitch were acquired during trial 10 and 90 in the Habituation and Acquisition phases allowing for early and late assessments in each phase. During the Acquisition phase, the first and third trials were CS+ trials to facilitate learning. All other trials were presented in a pseudorandomized order, such that no more than two CS+ trials would occur in sequence. All trials were separated by an inter-trial interval (ITI) randomly varying (uniform distribution) from 1.85 to 3.5 seconds in duration. All visual stimuli were presented using Psychtoolbox code (Brainard, 1997) on a Cambridge Research Systems Display ++ monitor (1,920 × 1,080 pixels, 120 Hz refresh rate), positioned approximately 120 cm in front of the participant.

Following data collection, a sequence of questions was used to assess how many participants differentiated the tones and if they learned the relationship between the CS+ and the US. Questions included how many distinct tones participants heard, an estimate of how many times they heard the US, and if there was a relationship between any of the sounds in the study as well as what that relationship was.

### Data Acquisition & Preprocessing

#### EEG Acquisition

EEG data were recorded at a 500 Hz sampling rate using an Electrical Geodesics (EGI) high input impedance system with a 128-channel (Ag-AgCl electrodes) HydroCel net, with impedances being kept below 60 kΩ. Online data were referenced to the vertex electrode (Cz), with a Butterworth filter applied during continuous recording. Following acquisition, EEG data were re-referenced to the grand average reference (i.e., averaged across all electrodes), and re-filtered with Butterworth low-pass (10th order, 3 dB point at 30 Hz) and high-pass (3rd order, 3 dB point at 1 Hz) filters. Next, continuous EEG data were separated into 3.6 second (1,801 sample points) trial epochs, spanning 600 ms (300 sample points) prior to tone onset and 3000 ms (1,500 sample points) post-onset.

Segmented trials were then evaluated for artifact contaminated trials using the Statistical Correction of Artifacts in Dense Array Studies (SCADS) procedure (Junghöfe et al., 2000). Eye artifacts were corrected with a regression-based EOG correction method (Schlögl et al., 2007, 2009) using electrodes above and below the eyes as well as electrode located at the outer canthi. Participants with a substantial number of trials lost due to artifacts (i.e., > 50% of all trials rejected) were excluded from further data analyses in both our primary (i.e., full behavioral and alpha-band power sample of 55) and secondary (i.e., reduced pupil sample of 35) data sets. The final sample had an average of 29.3 trials per pitch condition and phase (habituation, acquisition) retained. Importantly, a similar number of trials per condition within each phase were retained both our full behavioral and alpha-band power (Habituation: CS+ = 31; GS1 = 32; GS2 = 32; Acquisition: CS+ = 27; GS1 = 27; GS2 = 27).

#### Pupillometry Acquisition

Eye-tracking data were recorded using an EyeLink 1000 Plus eye tracker system with a 16 mm lens placed in front of the monitor, but outside of visible range for the participants’ lower visual field. The data were recorded at a 500 Hz sampling rate. Pupil diameter was quantified by fitting an ellipse to each participant’s pupil mass threshold. The system’s infrared signal illumination level was initially set to 100%, and individual adjustments were made for each participant based on their respective pupil and corneal reflection. Eye-tracking data were calibrated and validated using a nine-point grid. During this process, a white circle (1° visual angle) was presented at each of these nine points against a black background. As the participants’ eyes fixated on each location presentation, the lens and pupil thresholds were adjusted if pupil diameter was lost.

### Signal Processing

#### EEG analysis

Following artifact EEG preprocessing, a current source density (CSD) spatial transformation was applied to artifact-free trials, to increase spatial resolution. Using this method, signals originating from the cortical surfaces were estimated via the second spatial derivative (i.e., spatial Laplacian) from scalp topography. The equation proposed by Junghöfer and colleagues (1997) was used, suitable for dense-array EEG montages, with the recommended smoothing constant (*λ*) of 0.2. This CSD transformation heightens the interpretability of the topographical representation (Junghöfer et al., 1997; Kayser & Tenke, 2015) and thus assists in addressing questions regarding the cortical origin of conditioning-related power modulations in the alpha band.

Following the CSD spatial transformation, single-trial data from each participant, condition, and sensor were transformed into the time-frequency domain through convolution with a family of complex Morlet wavelets. These wavelets had center frequencies (*f)* between 2.50 and 27.48 Hz, spaced in even intervals of 0.278 Hz. Although the focus of this study was on alpha-band oscillations, frequencies in the entire range between 2.5 and 27.48 Hz were examined as recommended by current guidelines (Keil et al., 2022), to establish specificity and robustness of differences observed. A Morlet constant (*m*) of 10 was selected based to optimize the trade-off between temporal (*sigma_t*) and frequency, i.e., *sigma_f* = 1/(2 × π × *sigma_t*) smearing in the alpha range, given the ratio *m* = *f*/*sigma_f*, for every frequency in the analysis, *f*. At the center frequency of interest (10.27 Hz, the center of the canonical alpha band in young adults), this resulted in temporal and frequency smoothing values of 155 ms and 1.03 Hz, respectively. The absolute value from the convolution between the EEG data and the complex wavelets served as an estimate of time-varying power (Tallon-Baudry & Bertrand, 1999). Percent change in baseline alpha-band power for all EEG trials was calculated. The decision for this type of baseline adjustment was based on the presence of alpha-band activity in the baseline period, and because multiplicative changes in the alpha frequency range have been linked to meaningful differences in task performance (Foxe & Snyder, 2011; Jensen & Mazaheri, 2010). Specifically, the baseline period used for this correction was −422 to −202 ms prior to the onset of the stimulus, with this interval being selected to account for potential edge artifacts and the temporal smoothing value obtained from the wavelet transformation.

In addition to the trial-averaged time-frequency spectra, the trial-by-trial change in alpha power reduction was also examined in a post-hoc analysis, to assess how changes in oscillatory activity evolve as learning progresses. Baseline-corrected, single-trial time-by-frequency power at 10.27 Hz, recorded from nine mid-parietal sensors (site CPz and its nearest neighbors) during the time period of maximum alpha reduction (600-900ms post-stimulus) was averaged across electrodes and time points, for each participant. This window was chosen based on visual inspection of the grand mean time course of alpha-band changes. The resulting values (percent change in alpha power from baseline) were binned into 28 sequential groups of trials, 14 in Habituation and 14 in Acquisition, separately for each stimulus.

#### Pupil Quantification

Similar to EEG data, pupil diameter data were segmented into 3.6 second (1,801 sample points) epochs, with a 600 ms (300 sample points) before the tone and 3000 ms (1,500 sample points) after tone onset. Next, single-trials underwent artifact correction and detection procedures. First, a Butterworth low-pass filter (6^th^ order, 3dB point at 7.5 Hz) was applied to the data to attenuate large artifactual spikes. Then, two types of artifacts were addressed: 1) Time segments with missing values (e.g. due to blinks), 2) rapid pupil diameter changes exceeding 2.5 units change in pupil diameter from a previous sample point. These time segments were marked as artifacts and subsequently corrected using piece-wise cubic interpolation. Participants with inconsistent pupil tracking (defined as more than 50% interpolated data) were excluded from this analysis. As with the EEG data, the percentage of trial data interpolated did not significantly differ between pitch conditions in the Habituation (CS+ = 12.2%; GS1 = 14.1%; GS2 = 12.6%, *F* = 1.11, n.s.) or acquisition (CS+ = 17.1; GS1 = 15.9; GS2 = 15.9, *F* = 1.40, n.s.) phases in this reduced pupil data set. Finally, pupil data was baseline corrected as the percentage change from − 500 to 0 ms, relative to tone onset. Grand average pupil waveforms were computed across all pitch conditions in Habituation and Acquisition phases.

### Statistical Analyses

#### Overview

Affect ratings and physiological results were interpreted with the a-priori models of predicted outcomes: All-or-Nothing and Generalization. Each model had different weights reflecting the expected pattern of effects for the CS+, GS1, and GS2 trials. The All-or-Nothing model predicts a larger response only for the CS+ with no response for the GS1 or GS2 conditions; this pattern of effects is reflected in the weights of 2, –1, and –1 respectively. The Generalization model predicts the GS1 to evoke a similarly large response as the CS+ stimuli with weights of 1, 0.75, and –1.75. These weights were used to transform the valence and arousal ratings per person into a single value representing the model strength. For alpha-band power and pupil dilation, the model weights were used in the *F*-contrast permutation procedure. Both of these model fit analyses are described in the following sections.

#### Model fit analyses: Affective ratings

The ratings from each phase of the study were transformed into model fit scores by taking the inner product of the results by each of the three models’ weights. Thus, each participant had a model fit score for the All-or-Nothing and Generalization models for both arousal and valence ratings per the early and late stages of the Habituation and Acquisition phases. All model fit scores were tested for statistical significance via single-sample t-tests against a null hypothesis of zero. The critical *p*-value was set at Bonferroni corrected threshold of .003 found by taking the typical threshold of .05 divided by the total number of the 16 significance tests performed. The standardized mean difference (Cohen’s D) measure of effect size was reported with each t-test. For this measure, a value of 0.01 is considered very small, 0.2 is small, 0.5 is medium, 0.8 is large, 1.2 is very large, and 2.0 is thought of as huge (Cohen, 2013; Sawilowsky, 2009).

#### Model fit analyses: EEG alpha and pupil diameter

Both alpha-band power and pupil dilation were examined with general linear models using the model weights applied to the Acquisition phase of the study. To understand at which points model fits were statistically meaningful, permutation-controlled mass univariate tests were performed at each temporal and spatial data point (Blair & Karniski, 1993): *F*-contrasts for the three models were computed for each model fit producing a spatiotemporal map of each model’s *F*-values. This process was then repeated 5,000 times with each of the three conditions (i.e., CS+, GS1, and GS2) randomly permuted per participant. The maximum *F-*value for each electrode-by-time-by-frequency point permutation was used to create a non-parametric null distribution in which the 95^th^ quantile served as the critical *F*-value (Blair & Karniski, 1993; McTeague et al., 2015). For trial-averaged alpha-band power, this resulted in an *F_crit_* of 11.18. For trial-by-trial binned alpha changes across the experimental session, the *F_crit_* was 10.8. For pupil dilation, the maximum *F-*value per time point was used to form the null distribution with the same 95^th^ quantile, resulting in a critical *F*-value (*F_crit_*) of 9.3.

## Results

Post-experiment, the majority of participants reported the proper relationship between the CS+ and the US. Although only 15 out of 55 participants (27%) properly reported that there were 3 distinct tones, 52 (94.5%) reported there was a relationship between the tones and the noise. Furthermore, 43 (78%) correctly stated that either the lowest (21 out of 29; 72.4%) or highest pitch (22 out of 26; 84.6%) was associated with the US. Although 12 participants did not correctly state that the lowest or highest pitch was associated with the US, post-hoc analyses suggested they did not differ in their ratings or physiological reactivity compared to the rest of the sample (Supplemental Figures 1-2). They consistently rated the CS+ as more arousing and showed similar physiological reactivity to the CS+ as well.

### Arousal & Valence Ratings

The results for arousal and valence ratings can be seen in Figure 2. On a scale of 1 being completely unarousing and 1920 being completely arousing, the mean arousal rating in the early Habituation phase for the CS+ stimulus was M = 735, SE = 30.2; GS1 was M = 672, SE = 26.3; and GS2 was M = 733, SE = 29.6. For the late Habituation phase, CS+ was M = 687, SE = 34.3; GS1 was M = 704, SE = 29.5; and GS2 was M = 755, SE = 31.2. During the early Acquisition phase, the CS+ was M = 1210, SE = 38.2; GS1 was M = 746, SE = 29.4; and GS2 was M = 750, SE = 31.5. In the late Acquisition phase, the CS+ arousal rating was M = 1117, SE = 40.9; GS1 was M = 751, SE = 32.4; and GS2 was M = 737, SE = 31.5.

**Figure 2.**
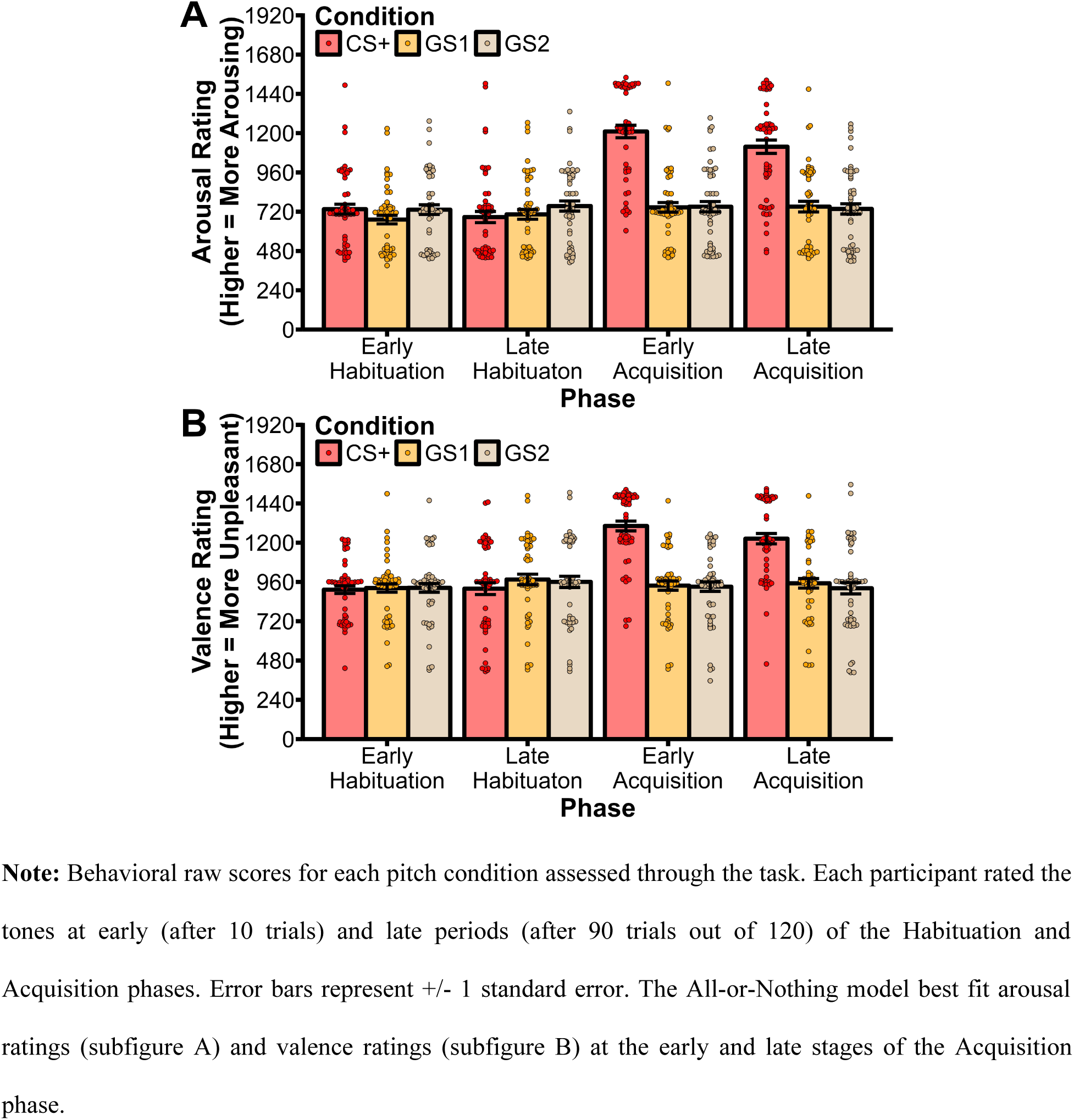
Arousal & Valence Raw Scores.

Valence ratings were on a similar scale as arousal self-reports, in which 1 was completely unpleasant and 1920 is completely pleasant. In the early Habituation phase, the CS+ stimuli wererated M = 913, SE = 23.5; GS1 was M = 923, SE = 25.1; and GS2 was M = 924, SE = 26.6. In the late Habituation phase, CS+ was M = 919, SE = 36.1; GS1 was M = 975, SE = 32.5; and GS2 was M = 960, SE = 34.4. For the early Acquisition phase, the CS+ was M = 1302, SE = 29.9; GS1 was M = 938, SE = 28.1; and GS2 was M = 931, SE = 28.6. In the late Acquisition phase, the CS+ arousal rating was M = 1224, SE = 31.7; GS1 was M = 952, SE = 29.1; and GS2 was M = 922, SE = 34.6.

### Arousal & Valence Model Fit Scores

As described in the methods section, arousal and valence ratings were analyzed by means of general linear models with weights defined by the three a-priori models for each phase of the study. The resulting fit scores were tested for significance via single-sample *t*-tests and a Bonferroni correct critical *p*-value of .003. The results of the *t*-tests can be seen in Table 1. The All-or-Nothing fit scores for arousal ratings were not significant in the early Habituation (M = 66.5, SE = 42.6) or late Habituation phases (M = −83.6, SE = 57.0). The arousal model fit scores were also not statistically significant for the Generalization model in both Habituation phases (early ratings M = −42.6, SE = 39.7; late ratings M = −106, SE = 49.1). In the Acquisition phases all model fit scores were statistically different with the largest effect sizes being for the All-or-Nothing model. In the early and late Acquisition phases respectively, the All-or-Nothing fit scores were M = 923, SE = 79.9 and M = 746, SE = 68.8, and the Generalization scores were M = 457, SE = 50.5 and M = 390, SE = 48.0.

**Table 1.**
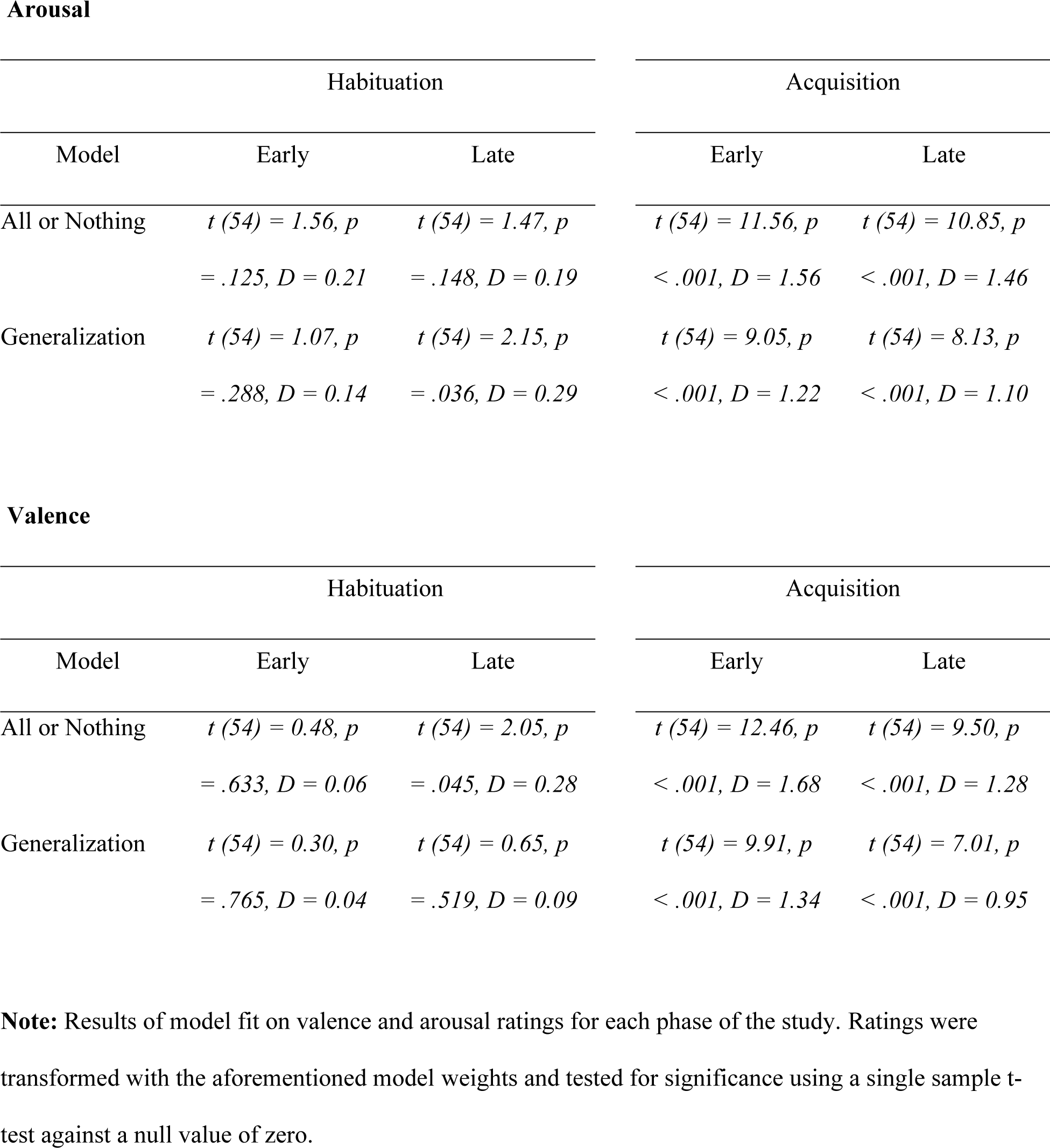
Statistical results of model fit scores.

For the valence ratings, model fit scores showed a similar pattern as the arousal ratings (Table 1). In the early and late Habituation phases, none of the model fit scores were statistically meaningful. In the Acquisition phases, all model fit scores were statistically significant with the All-or-Nothing transformed scores again featuring the largest effect sizes. In the early and late Habituation phases respectively, the model fit scores for the All-or-Nothing model were M = −20.6, SE = 42.8 and M = −97.1, SE = 47.4; for Generalization scores M = −11.70, SE = 39.1 and M = −30.3, SE = 46.7. In the early and late Acquisition phases, the All-or-nothing valence scores were M = 735, SE = 59.0 and M = 575, SE = 60.6; for Generalization scores M = 376, SE = 37.9 and M = 325, SE = 46.4.

### Alpha-Band Power

For the trial-averaged results, alpha-band power was found to be reduced during the presentation of each tone cue (Figure 3). It was primarily found over occipital-parietal areas even when baseline corrected (Figure 4). The *F*-contrast permutation procedure only found a significant alpha-band power reduction for the All-or-Nothing model. The *F*-values corresponding to that model exceeded the critical threshold at parieto-occipital electrode sites from 670 ms to 820 ms (Figure 5). This selective reduction in response to the conditioned threat cue was present in 40 out of the 55 participants (Figure 6).

**Figure 3.**
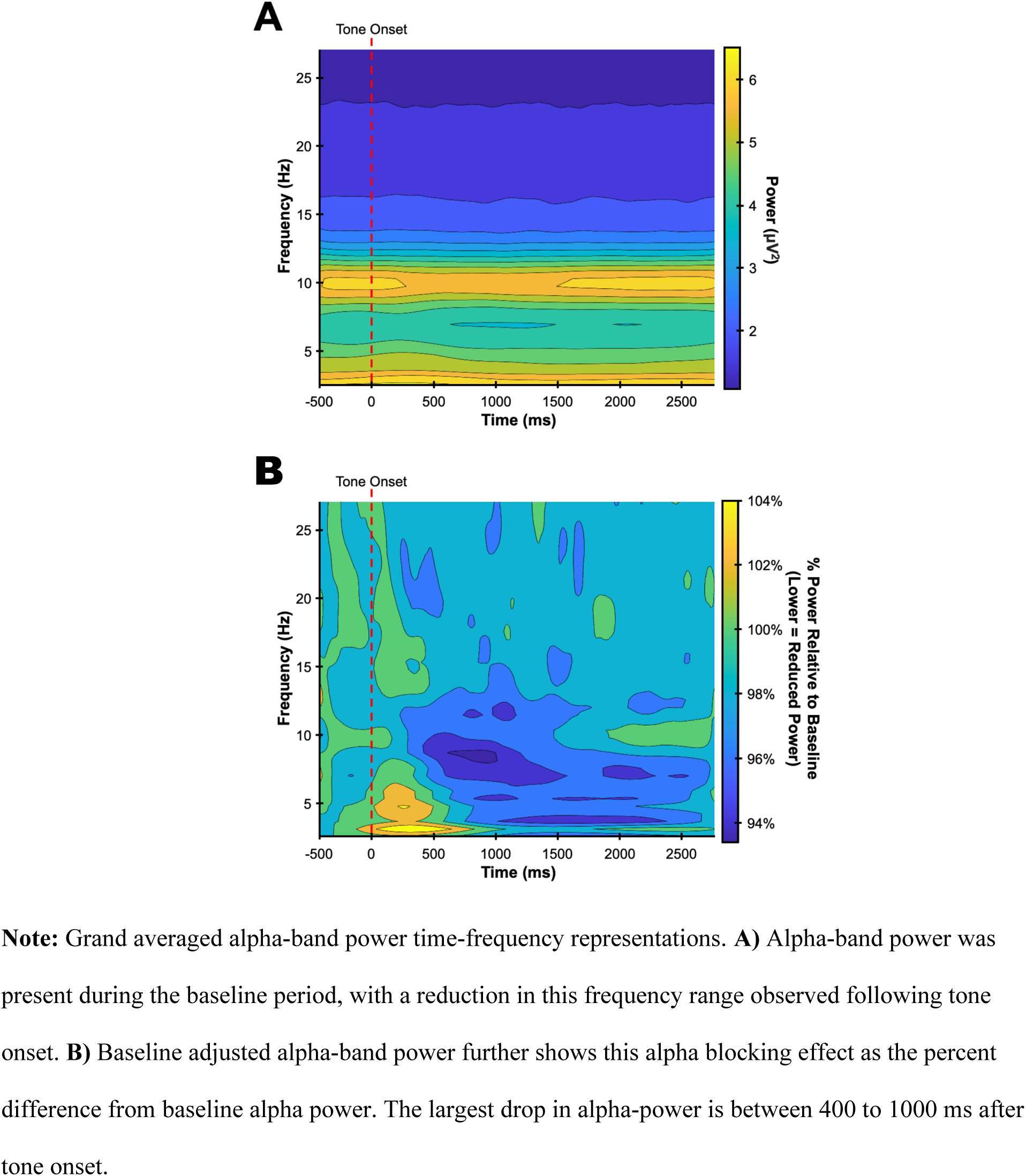
Time Frequency Alpha-Band Power.

**Figure 4.**
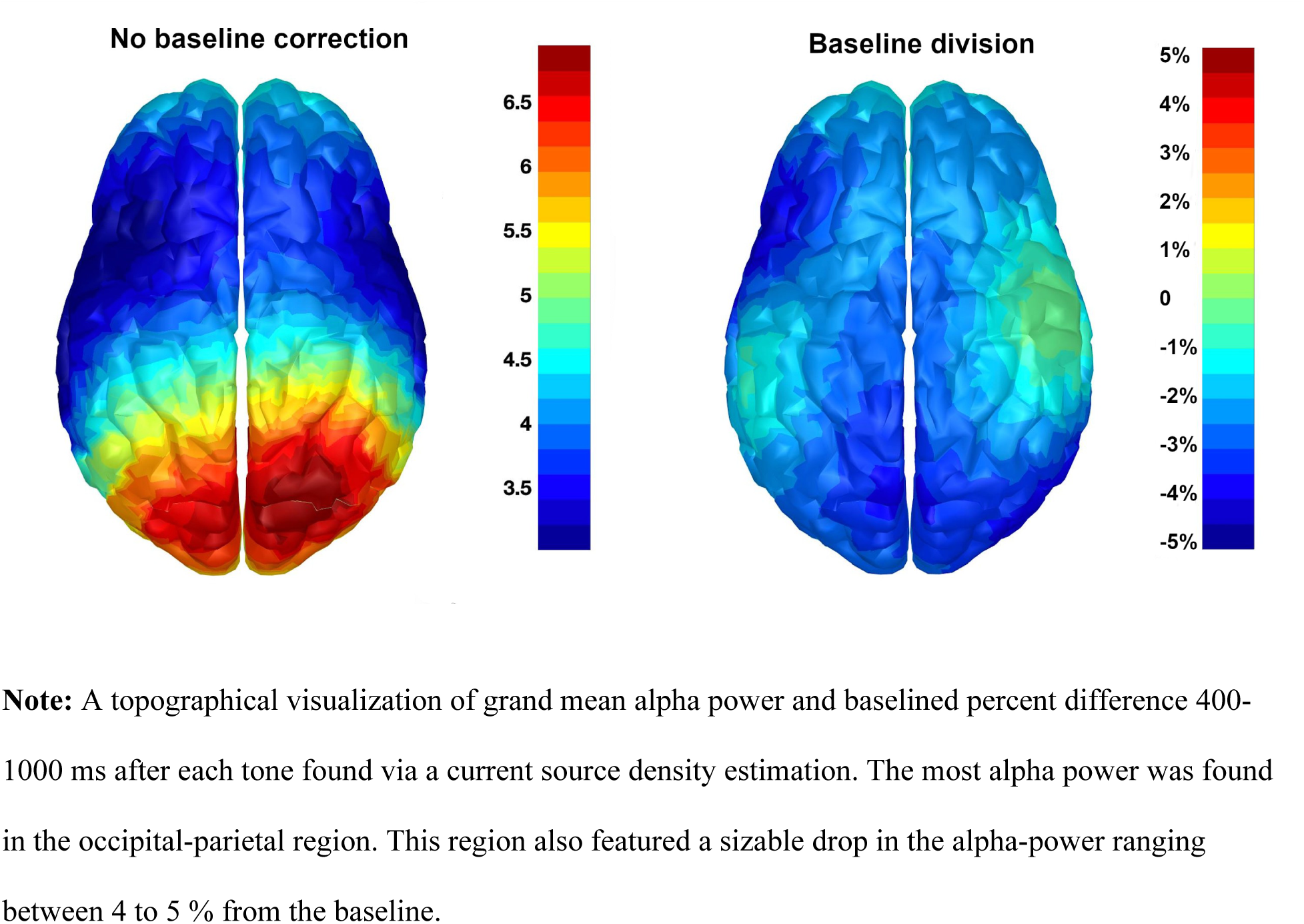
Grand Mean Alpha Power.

**Figure 5.**
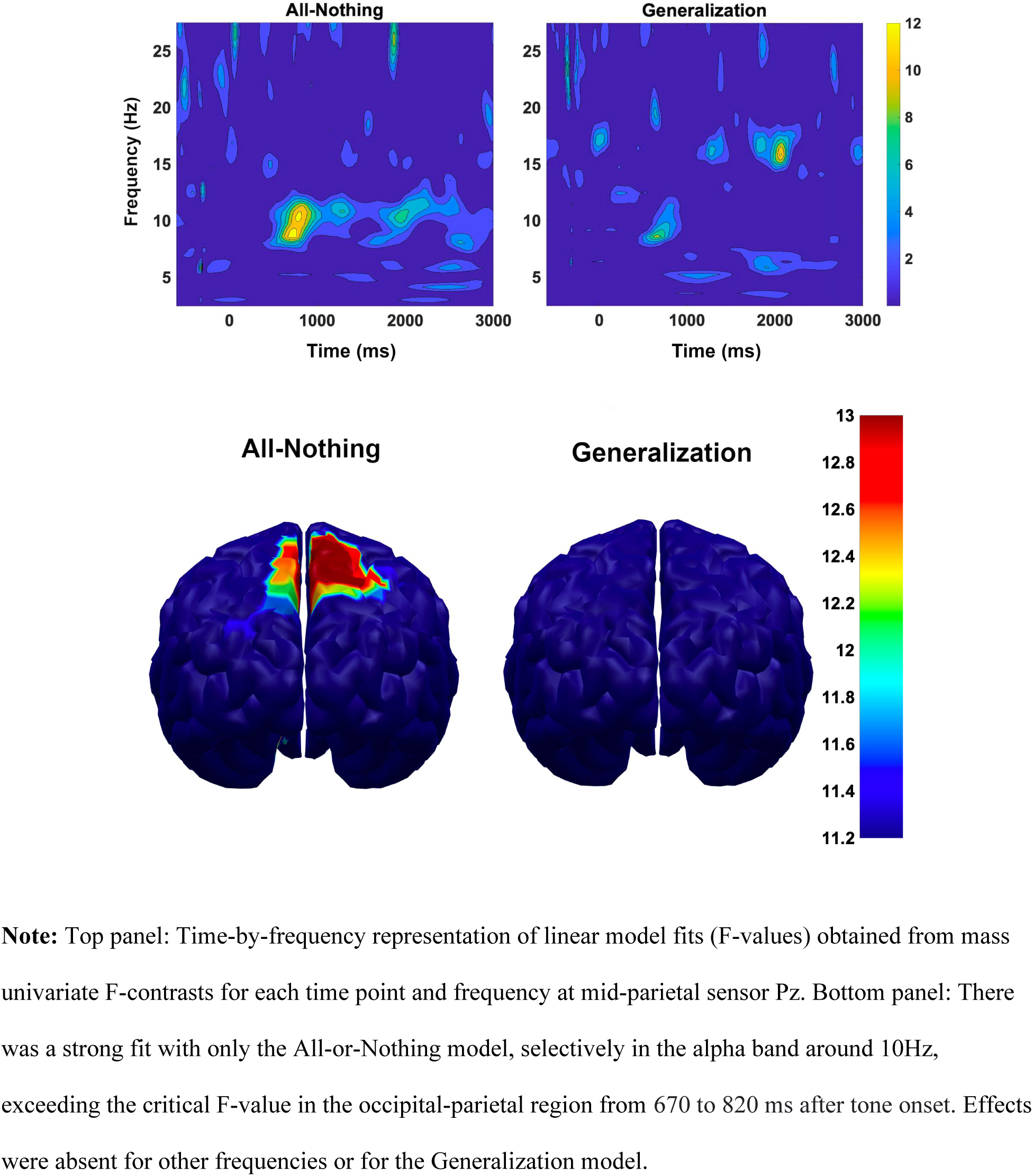
Model Fits for Alpha Band Power.

**Figure 6.**
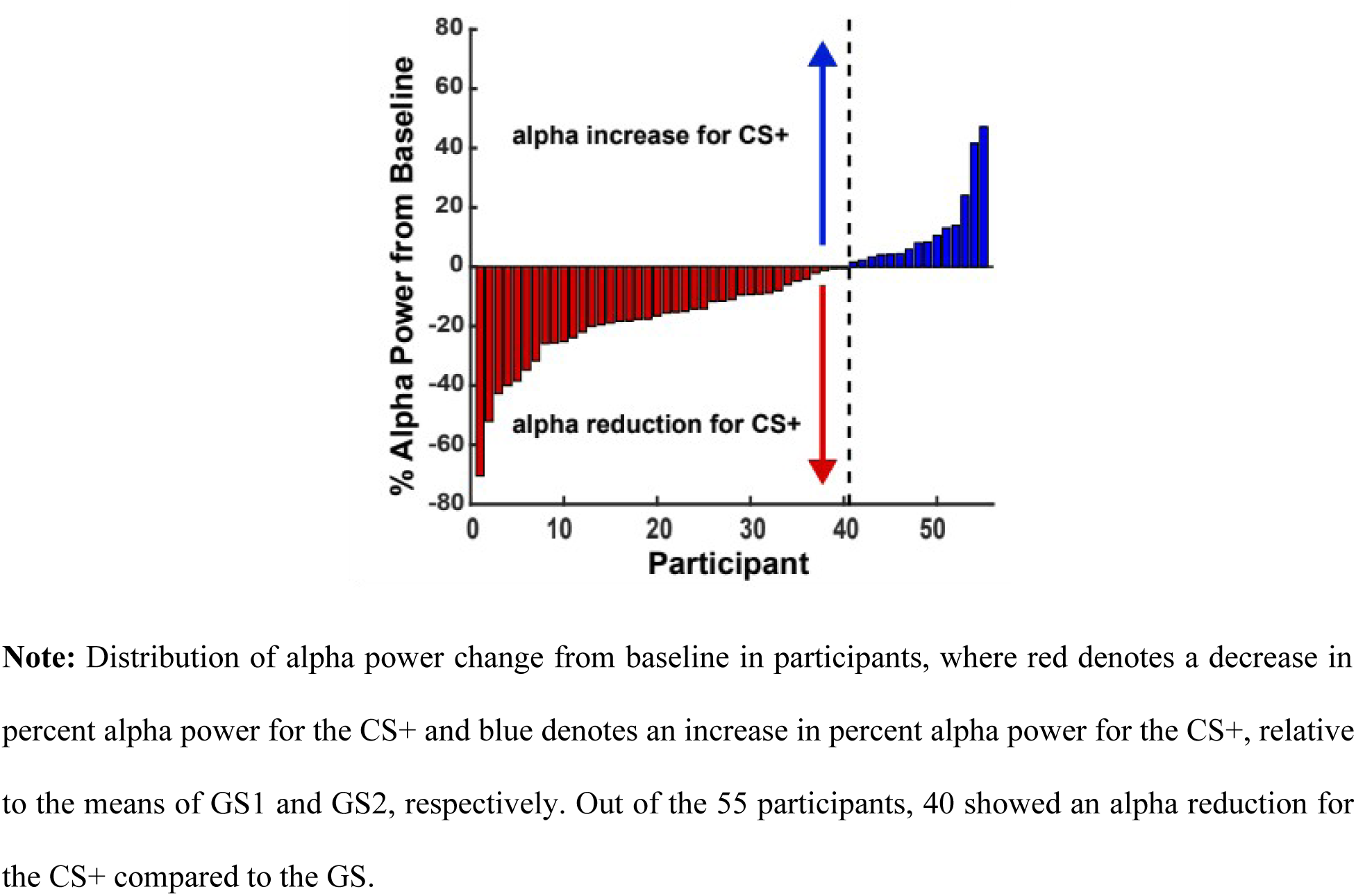
Distribution of % Alpha Power Change in Response to CS+.

When examining the trial-by-trial evolution of this selective alpha reduction across the experimental session (Figure 7), a significant *F*-contrast for the All-or-Nothing model was observed in trial bins 21-27, corresponding to the time between the 15^th^ and 32^nd^ trial of the Acquisition phase (out of 40 trials for each stimulus). Thus, selective alpha reduction in response to the conditioned threat cue appeared around mid-acquisition and lasted into the late phase of acquisition. Near the end of acquisition, the 27^th^ and 28^th^ trial bin no longer supported All-or-Nothing model and instead the data showed a generalization trend for the 27^th^ bin.

**Figure 7.**
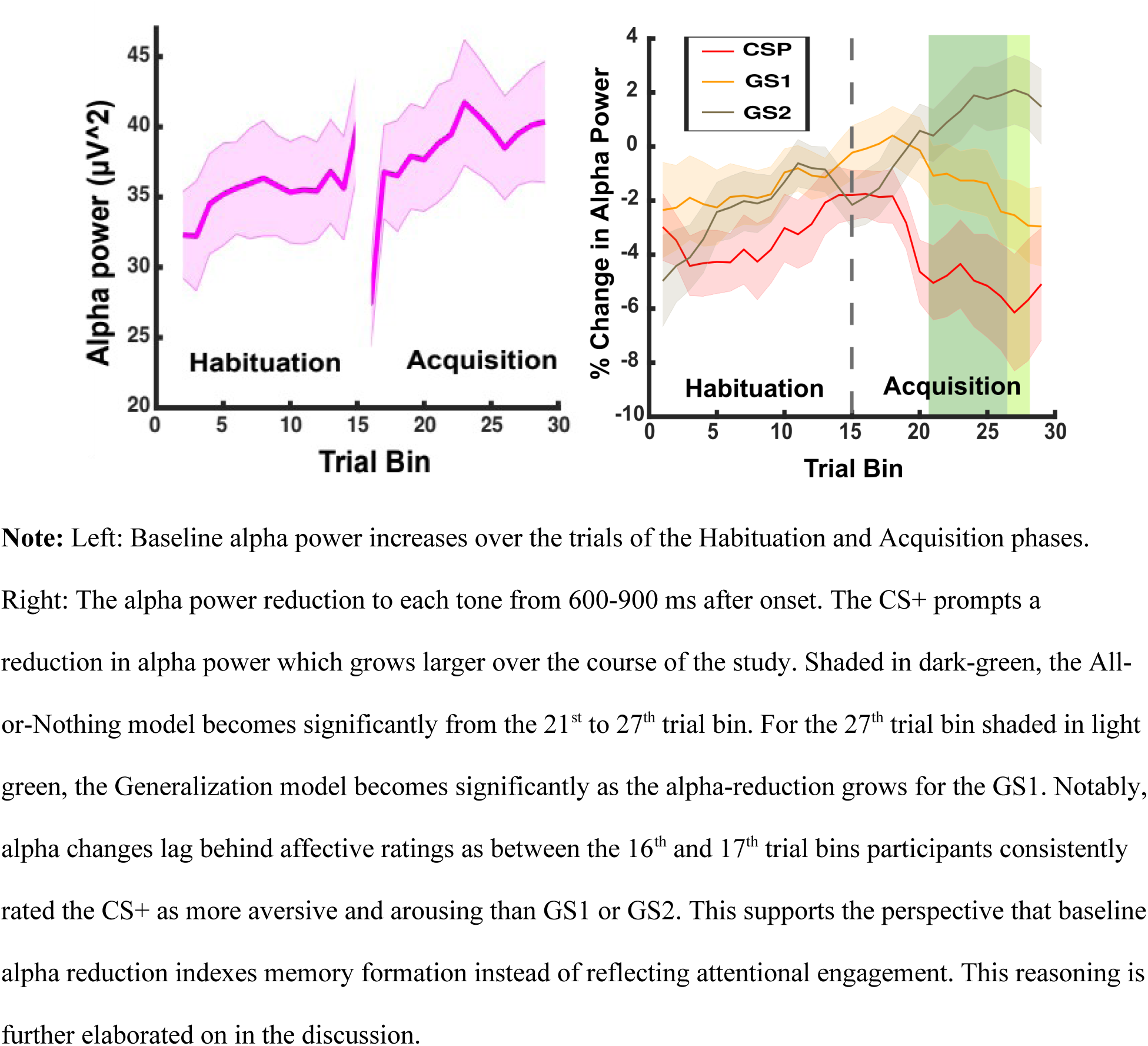
% Alpha Power Change Across Trials.

### Pupil Dilation

The average pupil waveforms by cue type are shown in Figure 8 for the Habituation and Acquisition phases. Consistent with the pattern visible in the grand mean waveforms, the permutation procedure showed that the only fit with the All-or-Nothing model surpassed the critical threshold at approximately 2500 ms (Figure 9). Paralleling alpha band power, the Generalization model did not surpass the critical threshold.

**Figure 8.**
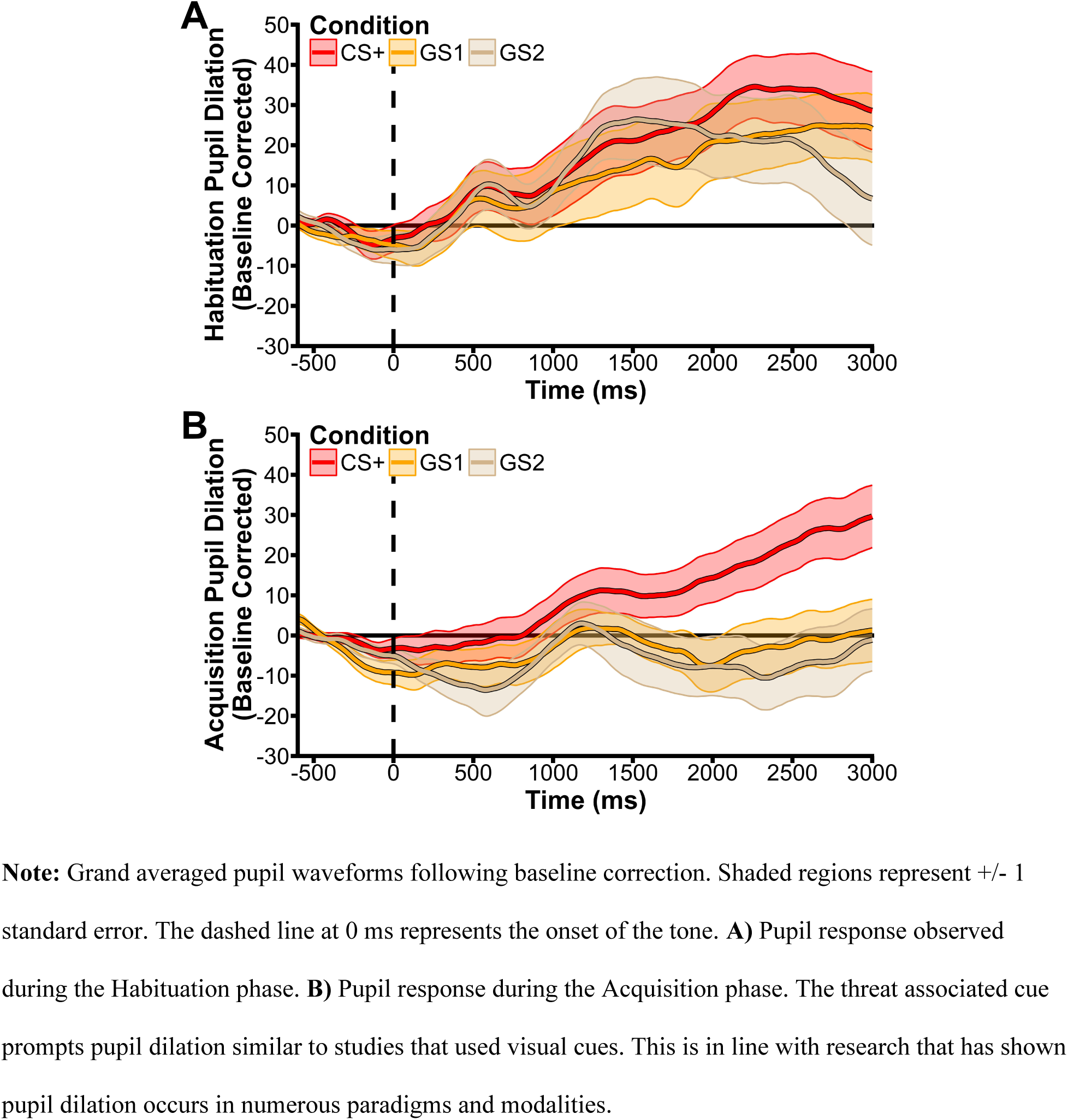
Grand Averaged Pupil Waveforms.

**Figure 9.**
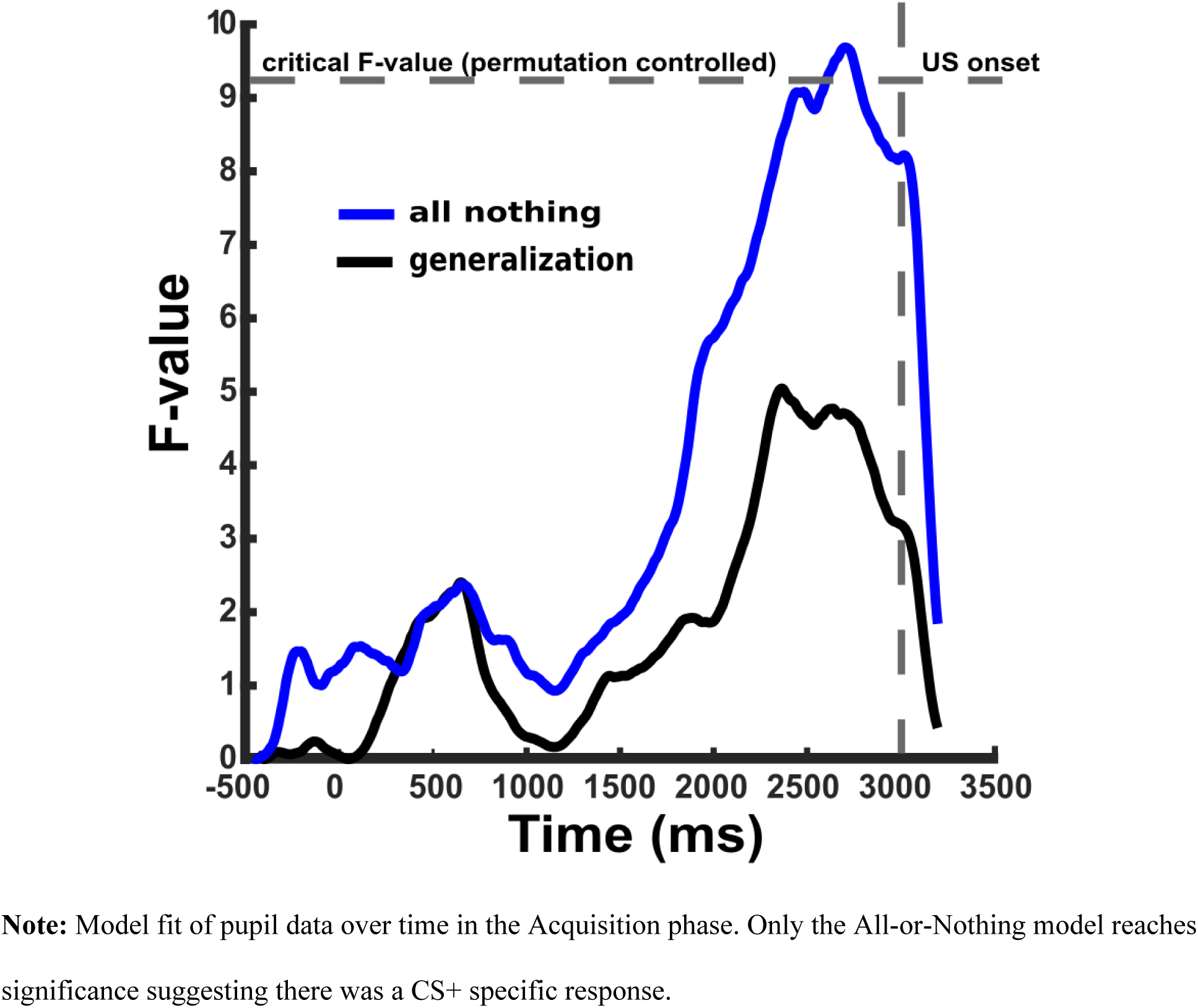
Model Fits for Pupil Timecourse.

## Discussion

In the present study, a differential aversive conditioning task was used to examine the hypothesis that alpha-band reductions serve as an index of ongoing cortical engagement with a conditioned auditory threat cue. Additionally, the study examined the extent to which alpha-power reduction in response to the CS+ generalizes to the most perceptually similar cue that was never paired to an aversive outcome (the GS1). The present study found strong support for the hypothesis that the selective reduction of parietal alpha oscillations reflects the engagement with conditioned auditory cues. The timing and topography of the alpha-band power reduction in the present study strongly resembled the occipital-parietal power changes observed with visual cues (Bacigalupo & Luck, 2022; Friedl & Keil, 2021; Yin et al., 2020), suggesting that suppression of alpha-band oscillation in parietal cortices serves as a supra-modal index of aversive conditioning. Unlike previous studies in the visual domain (Friedl & Keil, 2021), the present study did not show strong evidence of generalization effects. Among all variables, only trial-wise alpha power changes late in acquisition evinced generalization. This effect was driven by stronger alpha-power enhancement for the GS2 stimulus, compared to GS1, consistent with the overall trend of increased baseline alpha power during the experimental session, shown in Figure 7, left panel. This trend was accompanied by steadily increasing alpha power reduction throughout the acquisition session.

Against expectations, none of the other variables examined here—self-reported aversiveness and arousal, as well as pupil dilation—supported generalization. Accordingly, we found consistent support only for the All-or-Nothing model, when applied to ratings, pupil, and averaged parietal alpha power data. Future work will use smaller differences in pitch between CS+ and the GS, to examine the extent to which this absence of generalization reflects that the tones were too different.

At parietal sensor locations, 40 out of the 55 participants displayed a selectively larger reduction in alpha-power for the CS+, compared to the GS cues, showing that alpha reduction is a robust correlate of auditory conditioning. This effect for the CS+ emerged around mid-acquisition and lasted until the final trials of the experiment. This finding has several conceptual implications: Participants reported heightened displeasure and US expectancy for the CS+ already after 3 CS+ trials in the Acquisition phase, well before the emergence of selective alpha power reduction, which occurred after the 15^th^ CS+ trial. Thus, the present finding supports the notion that alpha reduction in response to conditioned threat reflects the formation of new aversive memory associations between the CS+ and the US (Yin et al., 2020). This is at odds with interpretation of selective alpha power reduction as reflective of attention to the CS+, a process that is expected to co-occur with the emergence of contingency awareness and self-reported expectancy (Bacigalupo & Luck, 2022). Providing further support to a memory formation account, post-hoc analysis of the GS stimuli in the present study shows that the difference between the safest (GS2) and the threat cue (CS+) remain robust until the end of the experimental session. In contrast, the more similar GS1 increasingly prompted alpha reduction as learning progress, consistent with generalization learning. Recent models of aversive conditioning, rooted in animal and human work, emphasize the steadily-changing neural dynamics as associative learning progresses (Li & Keil, 2023). Consistent with findings using fMRI (Yin et al., 2020) and event-related potentials (Thigpen, 2017), the present trial-by-trial analyses suggest that initial discrimination between threat and safety cues involves higher-order cortices involved in parietal alpha power changes. The observation that the discrimination between threat and safety cues is attenuated rather than increased later in the acquisition phase supports the notion that sustained aversive conditioning prompts discrimination at increasingly lower levels of the cortical hierarchy, eventually leading to threat representations in sensory areas (Li & Wilson, 2023).

Compared to previous research, the present results suggest that specific experimental conditions may facilitate auditory versus parietal alpha-band power changes. Hartmann and colleagues (2012) found an auditory cortex alpha reduction when participants thought they heard a “high” pitched tone which they were told predicted a noxious noise punishment; however, there was actually only one tone pitch presented to participants with pseudorandom feedback. In the present study, because the tones were mostly differentiable, perhaps the CS+ is quickly categorized and resources are diverted to processes related to the anticipation of the aversive US. In a preliminary report that included some of the current participants, primarily parietal alpha-power changes were also found (Ward et al., 2022). However, that study also included people with Misophonia, a disorder characterized by a heightened sensitivity to some noises.

Similar behavioral, pupil, and alpha-band power changes between visual and auditory conditioning paradigms are consistent with theoretical notions of how physiological changes are related to cognitive and behavioral processes. Physiological measures that index salience, attention, or emotion may occur as part of an adaptive coordinated response to motivationally relevant situations (Bradley, 2000; Lang et al., 1997). A threatening situation typically triggers concurrent physiological responses preparing the body to fight or flee, whether or not that action is carried out. So, if different paradigms evoke a similar action disposition, then similar physiological changes could be expected. For example, if it is presumed that pupil dilation can occur as a part of an adaptive general response whenever threat is anticipated, then it is not surprising that it is modulated by emotional scenes (Bradley et al., 2017), sounds (Partala & Surakka, 2003), emotional imagery (Henderson et al., 2018), and the anticipation of electrical shocks (Bitsios et al., 1996, 2004).

While generalization appears to emerge in the later trials of acquisition, it is unclear why the overall alpha-power reduction showed very little generalization to the GS1 when compared to visual-based studies (Friedl & Keil, 2021). Further experiments with different pitch gradients are likely necessary to examine the role of sensory distinctness in auditory generalization learning. There is considerable research on tone discrimination in humans (see, e.g., Moore, 2012). However, this work has not systematically been applied to aversive conditioning. Tone differentiation tasks typically involve presenting tones in quick succession and can differ widely based on the individual differences and features of the tone besides its frequency (Sek & Moore, 1995; Smith et al., 2017). These past studies suggest that participants should be able to differentiate tones much closer in pitch than the present research. However, during piloting for the present study, participants failed to correctly report CS-US contingencies when pitches were more proximal than in the present study. Interpreting pitch discrimination here is further complicated by the fact that the tones also contained a separate frequency aimed at evoking an auditory steady state response (not analyzed here) possibly making recognition more difficult. To clarify if generalization occurs, future research could either use tones more similar in pitch or add additional tones to see if this elicits a generalized response.

Better understanding alpha-power changes in differential aversive conditioning could have clinical significance in a variety of ways. Many psychiatric disorders are thought to feature an overgeneralization of fear responses, and this appears to be measurable for some conditions such as generalized anxiety disorders in differential conditioning paradigms (Lissek et al., 2010, 2014). However, as has been noted by others, auditory conditioning may sometimes be necessary as visual stimuli are not applicable for all participants such as children or those with disorders of consciousness (Kotchoubey, 2014; Rossetti & Laureys, 2015). Auditory conditioning paradigms could also provide unique insights into auditory specific symptoms. Noise sensitivity is associated with many psychiatric disorders (Stansfeld, 1992), particularly for the newly defined DSM disorder of Misophonia characterized by sensitive to loud and bodily noises. More generally, noise pollution itself appears to contribute to the prevalence of some mental health conditions (Hardoy et al., 2005; Zaman et al., 2022). Thus, future research to refine auditory conditioning paradigms and comparing the results against paired visual cues may lead to important new clinical findings.

In conclusion, baseline alpha-band power changes were studied in a differential aversive paradigm with auditory cues. Occipital-parietal alpha-power changes were found in most participants similar to visual cue studies, showing clear discrimination between conditioned threat and safety cues. There was very little evidence of generalization to the two other unpaired tones. Future work will carefully manipulate the psychophysical and aversive properties of tone cues to examine the relationship between sensory discriminability and aversive generalization. Overall, the present results are consistent with conceptual and mathematical models of aversive learning (Anderson, 2019; Miskovic & Keil, 2012), emphasizing that sustained aversive conditioning heightens sensory and attention changes, following a predictable time course (Li & Keil, 2023). Theoretical models and extant data suggest that aversive learning prompts initial increases in attentive processing of the CS+, followed by the emergence of sparse sensory representations of threat and safety features and a decrease in higher-order cortical involvement in threat processing. Future studies using mathematical modeling and multimodal imaging studies may refine and expand those models, ultimately informing a framework of aversive learning that includes specific cortical dynamics as a key mechanism.

## Acknowledgments

This research was funded by the Misophonia Research Fund by The Ream Foundation.

## Supplemental results

In the present study, participants reported after the EEG data-collection if there was a relationship between the tones and the US white noise burst. While 12 out of the 55 participants did not correctly state that either the highest or lowest tone was associated with the US, their results were not meaningfully different to affect the overall interpretations. The analyses and figures that led to that conclusion are presented here for interested readers.

### Ratings

A repeated measures ANOVA was used to assess if recognition of pairing had an interaction with the phase (Early Habituation, Late Habituation, Early Acquisition, and Late Acquisition) or condition (CS+, GS1, and GS2). The dependent variable for these ANOVAs was the difference in the CS+ rating from the GS1 and GS2. For arousal ratings the recognition by phase, recognition by condition, or recognition by condition by phase were not statistically significant; *F* (3, 159) = 0.72, *p =* .54, *F* (2, 106) = 0.25, *p =* .78, and *F* (6, 318) = 2.09, *p =* .05 respectively. This was also true for valence ratings; *F* (3, 159) = 1.51, *p =* .21, *F* (2, 106) = 2.00, *p =* .15, and *F* (6, 318) = 1.42, *p =* .20.

T-tests found that the participants that did not properly report the correct pairing still rated the CS+ as being more arousing than the GS1 and GS2 in Early Acquisition (*t* (11) = 8.89, *p* < .001) and Late Acquisition (*t* (11) = 5.61, *p* < .001). The CS+ was also rated as being more unpleasant for these participants in Early Acquisition (*t* (11) = 9.71, *p* < .001) and Late Acquisition (*t* (11) = 6.71, *p* < .001).

**Supplemental Figure 1.**
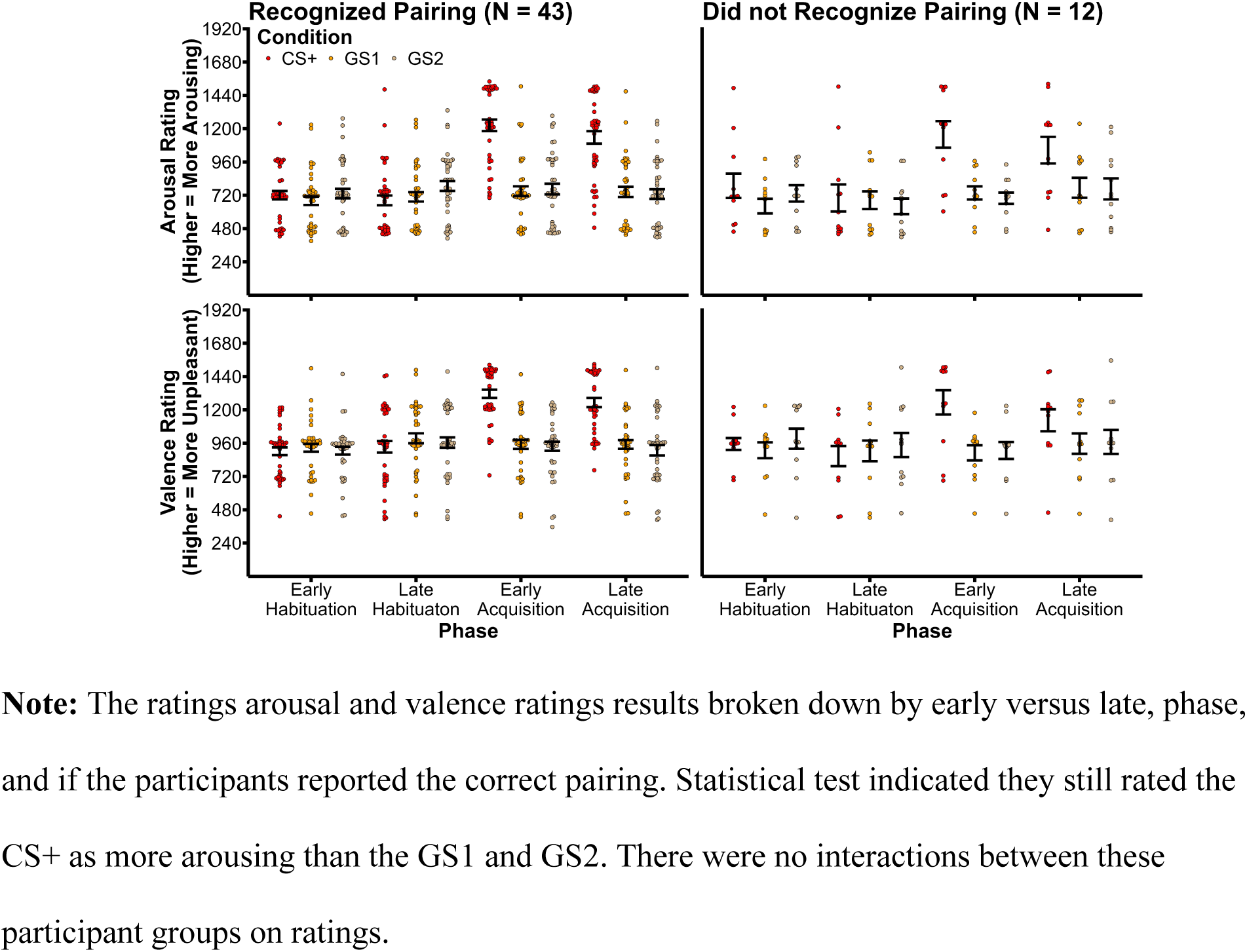
Ratings by phase, condition, and pairing recognition

### Alpha-band power

Here we assessed if the participants that reported the CS+ and US pairing incorrectly differed in alpha-band power changes compared to the result of the participants. A t-test found that overall alpha-band power change of the CS+ compared to the GS1 and GS2 was not different between those that reported the correct pairing or not; *t* (53) = 1.95, *p* = .067. Chi-squared test also found that the proportion of participants that had a heighten versus lower alpha-band power for the CS+ versus the GS stimuli was not different although the sample size is slightly too small for a credible estimate; χ2 (1) = .002, *p* = .96.

**Supplemental Figure 2.**
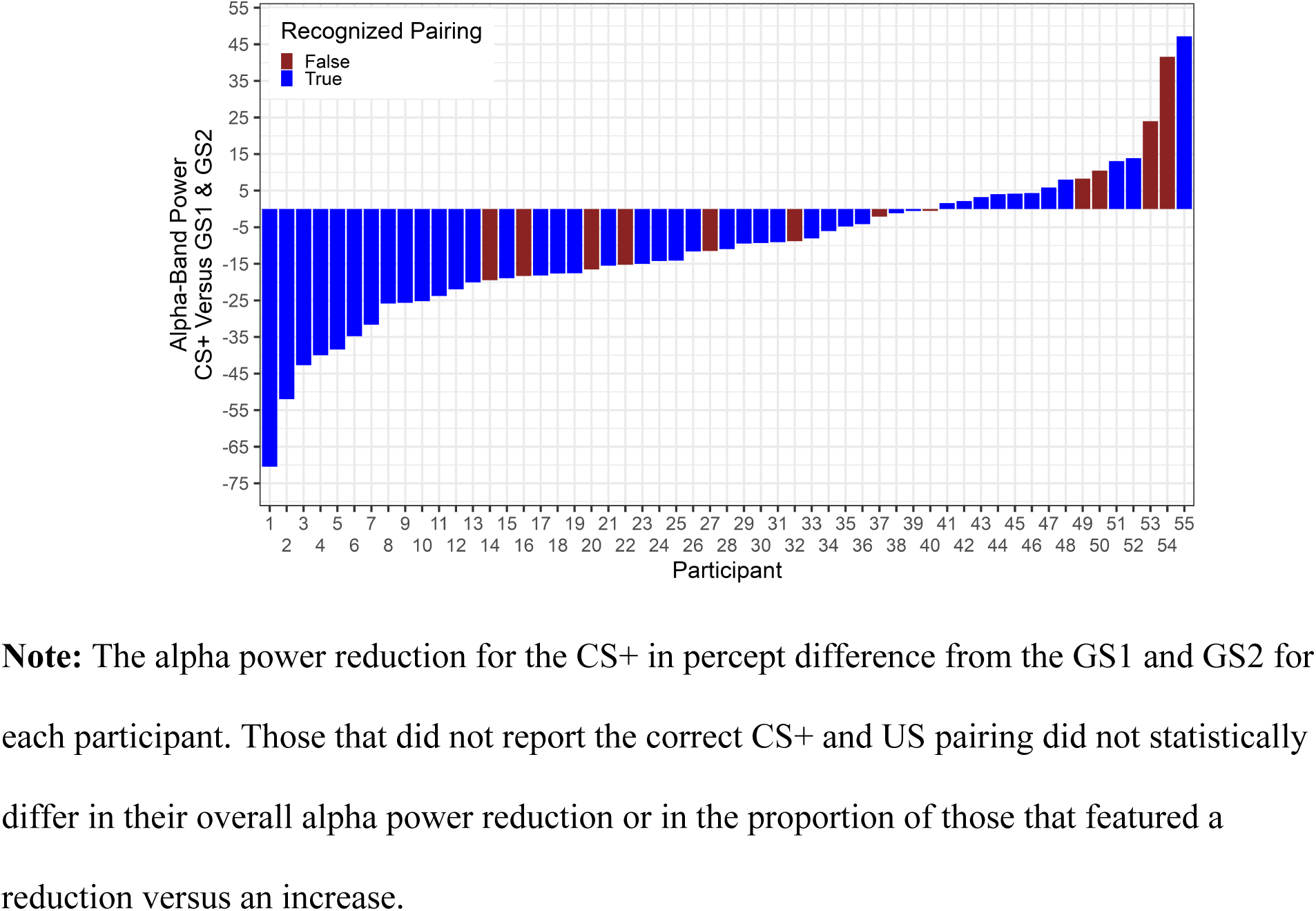
Alpha-power change for CS+ per participant.

## References

Anderson, B. A. (2019). Neurobiology of value-driven attention. Current Opinion in Psychology, 29, 27–33. 10.1016/j.copsyc.2018.11.004

Bacigalupo, F., & Luck, S. J. (2022). Alpha-band EEG suppression as a neural marker of sustained attentional engagement to conditioned threat stimuli. Social Cognitive and Affective Neuroscience, 17(12), 1101–1117. 10.1093/scan/nsac029

Bitsios, P., Szabadi, E., & Bradshaw, C. M. (1996). The inhibition of the pupillary light reflex by the threat of an electric shock: A potential laboratory model of human anxiety. Journal of Psychopharmacology, 10(4), 279–287. 10.1177/026988119601000404

Bitsios, P., Szabadi, E., & Bradshaw, C. M. (2004). The fear-inhibited light reflex: Importance of the anticipation of an aversive event. International Journal of Psychophysiology, 52(1), 87–95. 10.1016/j.ijpsycho.2003.12.006

Blair, R. C., & Karniski, W. (1993). An alternative method for significance testing of waveform difference potentials. Psychophysiology, 30(5), Article 5.

Bradley, M. M. (2000). Emotion and motivation. In J. T. Cacioppo, L. G. Tassinary, & G. Berntson (Eds.), Handbook of Psychophysiology (pp. 602–642). Cambridge University Press.

Bradley, M. M., & Lang, P. J. (1994). Measuring emotion: The Self-Assessment Manikin and the semantic differential. Journal of Behavior Therapy & Experimental Psychiatry, 25(1), Article 1.

Bradley, M. M., Sapigao, R. G., & Lang, P. J. (2017). Sympathetic ANS modulation of pupil diameter in emotional scene perception: Effects of hedonic content, brightness, and contrast. Psychophysiology, 54(10), 1419–1435. 10.1111/psyp.12890

Brainard, D. H. (1997). The Psychophysics Toolbox. Spat Vis, 10(4), Article 4.

Cohen, J. (2013). Statistical power analysis for the behavioral sciences. Academic press.

De Cesarei, A., & Codispoti, M. (2011). Affective modulation of the LPP and alpha-ERD during picture viewing. Psychophysiology, 48(10), Article 10. 10.1111/j.1469-8986.2011.01204.x

Ferrari, V., Mastria, S., & Codispoti, M. (2020). The interplay between attention and long-term memory in affective habituation. Psychophysiology, 57(6), e13572. 10.1111/psyp.13572

Foxe, J. J., & Snyder, A. C. (2011). The Role of Alpha-Band Brain Oscillations as a Sensory Suppression Mechanism during Selective Attention. Front Psychol, 2, 154. 10.3389/fpsyg.2011.00154

Friedl, W. M., & Keil, A. (2020). Effects of Experience on Spatial Frequency Tuning in the Visual System: Behavioral, Visuocortical, and Alpha-band Responses. Journal of Cognitive Neuroscience, 32(6), 1153–1169. 10.1162/jocn_a_01524

Friedl, W. M., & Keil, A. (2021). Aversive Conditioning of Spatial Position Sharpens Neural Population-Level Tuning in Visual Cortex and Selectively Alters Alpha-Band Activity. The Journal of Neuroscience, 41(26), 5723–5733. 10.1523/JNEUROSCI.2889-20.2021

Hardoy, M. C., Carta, M. G., Marci, A. R., Carbone, F., Cadeddu, M., Kovess, V., Dell’Osso, L., & Carpiniello, B. (2005). Exposure to aircraft noise and risk of psychiatric disorders: The Elmas survey: Aircraft noise and psychiatric disorders. Social Psychiatry and Psychiatric Epidemiology, 40(1), 24–26. 10.1007/s00127-005-0837-x

Hartmann, T., Schlee, W., & Weisz, N. (2012). It’s only in your head: Expectancy of aversive auditory stimulation modulates stimulus-induced auditory cortical alpha desynchronization. NeuroImage, 60(1), 170–178. 10.1016/j.neuroimage.2011.12.034

Henderson, R. R., Bradley, M. M., & Lang, P. J. (2018). Emotional imagery and pupil diameter. Psychophysiology, 55(6), e13050. 10.1111/psyp.13050

Jensen, O., & Mazaheri, A. (2010). Shaping functional architecture by oscillatory alpha activity: Gating by inhibition. Front Hum Neurosci, 4, 186. 10.3389/fnhum.2010.00186

Junghöfe, M., Elbert, T., Tucker, D. M., & Rockstroh, B. (2000). Statistical control of artifacts in dense array EEG/MEG studies. Psychophysiology, 37(4), Article 4.

Junghöfer, M., Elbert, T., Leiderer, P., Berg, P., & Rockstroh, B. (1997). Mapping EEG-potentials on the surface of the brain: A strategy for uncovering cortical sources. Brain Topogr, 9(3), Article 3.

Kamin, L. J. (1956). The effects of termination of the CS and avoidance of the US on avoidance learning. Journal of Comparative and Physiological Psychology, 49(4), 420–424. 10.1037/h0088011

Kayser, J., & Tenke, C. E. (2015). On the benefits of using surface Laplacian (current source density) methodology in electrophysiology. International Journal of Psychophysiology, 97(3), 171–173. 10.1016/j.ijpsycho.2015.06.001

Keil, A., Bernat, E. M., Cohen, M. X., Ding, M., Fabiani, M., Gratton, G., Kappenman, E. S., Maris, E., Mathewson, K. E., Ward, R. T., & Weisz, N. (2022). Recommendations and publication guidelines for studies using frequency domain and time-frequency domain analyses of neural time series. Psychophysiology, 59(5), e14052. 10.1111/psyp.14052

Kotchoubey, B. (2014). Event-related potentials in disorders of consciousness. In Clinical Neurophysiology in Disorders of Consciousness: Brain Function Monitoring in the ICU and Beyond (pp. 107–123). Springer.

Lang, P. J., Bradley, M. M., & Cuthbert, B. N. (1997). Motivated Attention: Affect, Activation, and Action. In P. J. Lang, R. F. Simons, & M. T. Balaban (Eds.), Attention and Orienting: Sensory and Motivational Processes (pp. 97–135). Lawrence Erlbaum Associates.

Li, W., & Keil, A. (2023). Sensing fear: Fast and precise threat evaluation in human sensory cortex. Trends in Cognitive Sciences, 27(4), 341–352. 10.1016/j.tics.2023.01.001

Li, W., & Wilson, D. A. (2023). Threat Memory in the Sensory Cortex: Insights from Olfaction. The Neuroscientist, 10738584221148994. 10.1177/10738584221148994

Lissek, S., Bradford, D. E., Alvarez, R. P., Burton, P., Espensen-Sturges, T., Reynolds, R. C., & Grillon, C. (2014). Neural substrates of classically conditioned fear-generalization in humans: A parametric fMRI study. Soc Cogn Affect Neurosci, 9(8), Article 8. 10.1093/scan/nst096

Lissek, S., Rabin, S., Heller, R. E., Lukenbaugh, D., Geraci, M., Pine, D. S., & Grillon, C. (2010). Overgeneralization of conditioned fear as a pathogenic marker of panic disorder. Am J Psychiatry, 167(1), Article 1. 10.1176/appi.ajp.2009.09030410

McTeague, L. M., Gruss, L. F., & Keil, A. (2015). Aversive learning shapes neuronal orientation tuning in human visual cortex. Nature Communications, 6, 7823. 10.1038/ncomms8823

Miskovic, V., & Keil, A. (2012). Acquired fears reflected in cortical sensory processing: A review of electrophysiological studies of human classical conditioning. Psychophysiology, 49(9), Article 9. 10.1111/j.1469-8986.2012.01398.x

Moore, B. C. (2012). An introduction to the psychology of hearing. Brill.

Panitz, C., Keil, A., & Mueller, E. M. (2019). Extinction-resistant attention to long-term conditioned threat is indexed by selective visuocortical alpha suppression in humans. bioRxiv, 533141. 10.1101/533141

Partala, T., & Surakka, V. (2003). Pupil size variation as an indication of affective processing. International Journal of Human-Computer Studies, 59(1–2), 185–198. 10.1016/S1071-5819(03)00017-X

Pavlov, Y. G., & Kotchoubey, B. (2019). Classical conditioning in oddball paradigm: A comparison between aversive and name conditioning. Psychophysiology, 56(7), e13370. 10.1111/psyp.13370

Rosenthal, R., & Rosnow, R. L. (1985). Contrast Analysis: Focused Comparisons in the Analysis of Variance. CUP Archive.

Rossetti, A. O., & Laureys, S. (Eds.). (2015). Clinical Neurophysiology in Disorders of Consciousness: Brain Function Monitoring in the ICU and Beyond. Springer Vienna. 10.1007/978-3-7091-1634-0

Sawilowsky, S. S. (2009). New Effect Size Rules of Thumb. Journal of Modern Applied Statistical Methods, 8(2), 597–599. 10.22237/jmasm/1257035100

Schlögl, A., Keinrath, C., Zimmermann, D., Scherer, R., Leeb, R., & Pfurtscheller, G. (2007). A fully automated correction method of EOG artifacts in EEG recordings. Clinical Neurophysiology, 118(1), 98–104. 10.1016/j.clinph.2006.09.003

Schlögl, A., Ziehe, A., & Müller, K.-R. (2009). Automated ocular artifact removal: Comparing regression and component-based methods. Nature Precedings. 10.1038/npre.2009.3446.1

Sek, A., & Moore, B. C. J. (1995). Frequency discrimination as a function of frequency, measured in several ways. The Journal of the Acoustical Society of America, 97(4), 2479–2486. 10.1121/1.411968

Smith, L. M., Bartholomew, A. J., Burnham, L. E., Tillmann, B., & Cirulli, E. T. (2017). Factors affecting pitch discrimination performance in a cohort of extensively phenotyped healthy volunteers. Scientific Reports, 7(1), 16480. 10.1038/s41598-017-16526-8

Sperl, M. F. J., Panitz, C., Hermann, C., & Mueller, E. M. (2016). A pragmatic comparison of noise burst and electric shock unconditioned stimuli for fear conditioning research with many trials. Psychophysiology, 53(9), 1352–1365. 10.1111/psyp.12677

Stansfeld, S. A. (1992). Noise, noise sensitivity and psychiatric disorder: Epidemiological and psychophysiological studies. Psychological Medicine. Monograph Supplement, 22, 1–44. 10.1017/S0264180100001119

Tallon-Baudry, C., & Bertrand, O. (1999). Oscillatory gamma activity in humans and its role in object representation. Trends in Cognitive Sciences, 3(4), Article 4.

Thigpen, N. N. (2017). The malleability of emotional perception: Short-term plasticity in retinotopic neurons accompanies the formation of perceptual biases to threat. Journal of Experimental Psychology: General, 146(4), 464. 10.1037/xge0000283

Ward, R. T., Gilbert, F. E., Pouliot, J., Chiasson, P., McIlvanie, S., Traiser, C., Riels, K., Mears, R., & Keil, A. (2022). The Relationship Between Self-Reported Misophonia Symptoms and Auditory Aversive Generalization Leaning: A Preliminary Report. Frontiers in Neuroscience, 16, 899476. 10.3389/fnins.2022.899476

Wu, M. S., Lewin, A. B., Murphy, T. K., & Storch, E. A. (2014). Misophonia: Incidence, Phenomenology, and Clinical Correlates in an Undergraduate Student Sample. Journal of Clinical Psychology, 70(10), Article 10. 10.1002/jclp.22098

Yin, S., Bo, K., Liu, Y., Thigpen, N., Keil, A., & Ding, M. (2020). Fear conditioning prompts sparser representations of conditioned threat in primary visual cortex. Social Cognitive and Affective Neuroscience, 15(9), Article 9. 10.1093/scan/nsaa122

Zaman, M., Muslim, M., & Jehangir, A. (2022). Environmental noise-induced cardiovascular, metabolic and mental health disorders: A brief review. Environmental Science and Pollution Research, 29(51), 76485–76500. 10.1007/s11356-022-22351-y

